# The transfer function of the rhesus macaque oculomotor system for small-amplitude slow motion trajectories

**DOI:** 10.1101/359836

**Authors:** Julianne Skinner, Antimo Buonocore, Ziad M. Hafed

## Abstract

Two main types of small eye movements occur during gaze fixation: microsaccades and slow ocular drifts. While microsaccade generation has been relatively well-studied, ocular drift control mechanisms are unknown. Here we explored the degree to which monkey smooth eye movements, on the velocity scale of slow ocular drifts, can be generated systematically. Two male rhesus macaque monkeys tracked a spot moving sinusoidally, but slowly, along the horizontal or vertical directions. Maximum target displacement in the motion trajectory was 30 min arc (0.5 deg), and we varied the temporal frequency of target motion from 0.2 to 5 Hz. We obtained an oculomotor “transfer function” by measuring smooth eye velocity gain (relative to target velocity) as a function of frequency, similar to past work with large-amplitude pursuit. Monkey eye velocities as slow as those observed during slow ocular drifts were clearly target-motion driven. Moreover, like with large-amplitude smooth pursuit, eye velocity gain varied with temporal frequency. However, unlike with large-amplitude pursuit, exhibiting low-pass behavior, small-amplitude motion tracking was band-pass with the best ocular movement gain occurring at ~0.8-1 Hz. When oblique directions were tested, we found that the horizontal component of pursuit gain was larger than the vertical component. Our results provide a catalogue of the control abilities of the monkey oculomotor system for slow target motions, and they also support the notion that smooth fixational ocular drifts are controllable. This has implications for neural investigations of drift control and the image-motion consequences of drifts on visual coding in early visual areas.

## Introduction

Small fixational eye movements continuously move the retinal image, even though they never deviate the center of gaze (or fovea) away from objects of interest. Such continuous movements mean that the neural signals representing input images in a variety of early visual areas are continuously modulated, and this has implications for theoretical interpretations of neural variability across trials, as well as population correlations in simultaneously recorded neurons (McFarland et al. 2016; Rucci and Victor 2015). Therefore, understanding the neural control mechanisms for fixational eye movements is an important first step towards understanding the full impact of these eye movements on neural coding, and ultimately visual perception.

Fixational eye movements come in two main flavors: microsaccades, which are rapid gaze shifts that look like big saccades; and smooth ocular drifts, which are slow changes in eye position occurring in between microsaccades (Barlow 1952; Murphy et al. 1975; Nachmias 1961). Even though microsaccades have received a substantial amount of research attention recently (Hafed 2011; Hafed et al. 2015; Krauzlis et al. 2017; Rolfs 2009), drifts remain to be relatively underexplored. For example, the mechanisms for generating ocular drifts are unknown, whereas microsaccade generation has been studied in a variety of different brain areas (Arnstein et al. 2015; Hafed et al. 2009; Hafed and Krauzlis 2012; Peel et al. 2016; Sun et al. 2016; Van Horn and Cullen 2012). Moreover, uncovering detailed neural mechanisms for microsaccade generation in certain key brain areas was instrumental for uncovering important, previously unappreciated, consequences of these eye movements on vision, perception, and even cognition (Chen et al. 2015; Hafed 2013; 2011; Hafed et al. 2015; Krauzlis et al. 2017; Tian et al. 2016; Veale et al. 2017). Thus, exploring the neural mechanisms for slow ocular drift generation is a worthwhile effort, especially given the fact that during fixation, microsaccades are brief events in an otherwise continuous sea of ocular drifts.

Here, towards approaching that ultimate goal, we aimed to characterize the degree to which the control system for eye position in the macaque monkey brain is able to generate very slow smooth pursuit eye movements, on the velocity scale of fixational ocular drifts. In other words, is it possible for monkeys to volitionally generate a slow eye movement that is as small in amplitude and velocity as slow ocular drifts, but that is clearly controlled and with a predictable motion trajectory? We designed behavioral experiments motivated by potential analogies that one can make between slow ocular drifts and smooth pursuit eye movements (Cunitz 1970; Martins et al. 1985; Nachmias 1961), similar to analogies that one makes between microsaccades and larger saccades (Hafed 2011; Krauzlis et al. 2017). After all, smooth pursuit eye movements are (relatively) slow rotations of the eyeball, similar in nature to slow ocular drifts, and the circuits for generating such smooth pursuit eye movements are well studied (Krauzlis 2004). Moreover, in general, the functional goals of smooth pursuit are primarily to: (1) stabilize retinal image motions of a stimulus, and (2) optimize the position of the eye such that the stimulus is within the fovea. Both of these goals are also the same for gaze fixation, even in the absence of microsaccades.

We presented macaque monkeys with small-amplitude sinusoidal target trajectories, and we asked how well their small-amplitude slow eye movements can track such trajectories. Our approach was to assume that for the frequency ranges that we studied, the oculomotor system may behave, to a first approximation, like a linear system. This means that we can present a single frequency and measure the response, and then test another frequency, and so on. The gain and phase lag of tracking at each frequency can thus allow estimating an “equivalent” transfer function of the oculomotor system (Ohashi and Mizukoshi 1991). Previous attempts like this with larger-amplitude (and faster) eye movements in smooth pursuit were effective, and showed that pursuit tracks very well for low frequencies (<1 Hz), but that it then behaves relatively poorly with higher frequencies (exhibiting both lower gain and larger phase lag) (Bahill and McDonald 1983; Collewijn and Tamminga 1984; Fabisch et al. 2009; Martins et al. 1985; Ohashi and Mizukoshi 1991; Rottach et al. 1996). We were interested in what happens in the monkey with much smaller and slower eye movements. The key comparison was to see whether controlled slow eye movements as slow as fixational ocular drifts would be possible. Critically, we performed our experiments on monkeys in order to demonstrate that these animals are instrumental for ultimately uncovering the neural control mechanisms for slow ocular drifts, and also in order to complement earlier human work on the topic (Cunitz 1970; de Bie and van den Brink 1986; Martins et al. 1985; Murphy et al. 1975; Nachmias 1961; Wyatt and Pola 1981).

## Materials and Methods

### Animal preparation and laboratory setup

We recorded eye movements from two male rhesus macaque (macaca mulatta) monkeys (monkey A and monkey M) aged 6-7 years. We implanted one eye (left for monkey A and right for monkey M) in each animal with a scleral search coil for eye tracking using the magnetic induction technique (Fuchs and Robinson 1966; Judge et al. 1980). The magnetic induction and eye movement measurement system that we used was provided by Remmel Labs, Texas, USA, and its technical specifications are described in (Remmel 1984). Our implanted coils were made out of steel wire obtained from Baird Industries, New Jersey, USA (part number: 300ft S170012a7-fep). We made four loops of wire in each coil to boost the signal and minimize noise. Surgical procedures for implantation were similar to those described earlier (Chen and Hafed 2013; Hafed and Ignashchenkova 2013), and we implanted the coils around the eye orbit anterior to the extra-ocular muscle insertions). The experiments were approved by ethics committees at the regional governmental offices of the city of Tübingen, and they were in accordance with European Union guidelines on animal research, as well as the associated implementations of these guidelines in German law.

We used monkeys in this study for two important reasons. First, the monkeys had scleral search coils implanted, which allowed the most precise measurement of small-amplitude slow eye movements, including fixational drifts. Video-based eye trackers (used in most human studies) are less reliable than scleral search coils for measuring slow eye movements (Chen and Hafed 2013; Choe et al. 2016; Kimmel et al. 2012). Second, and more importantly, the monkeys are now being used in neurophysiological recording experiments, such that direct neural correlates of our observations here can be identified and disseminated; particularly to complement earlier human work on similar questions (Cunitz 1970; de Bie and van den Brink 1986; Martins et al. 1985; Murphy et al. 1975; Nachmias 1961).

During data collection, the animals were seated in a primate chair 73 cm from a linearized (i.e. calibrated) CRT computer monitor (ViewSonic, model PF817, manufactured in 2000) in an otherwise dark room. The monitor had a pixel resolution of 34 pixels/deg and a refresh rate of 120 Hz. Stimuli were presented over a uniform gray background (29.7 Cd/m^2^ in Experiment 1, and either 29.7 or 4.4 Cd/m^2^ in Experiment 2). A small white spot (~5 x 5 min arc square) having 86 Cd/m^2^ luminance in Experiment 1 and either 86 or 48.1 Cd/m^2^ in Experiment 2 was used as the moving target for smooth pursuit eye movements (see *Behavioral tasks* below).

Graphics on the CRT monitor were presented by a computer running Matlab’s Psychophysics Toolbox (Brainard 1997; Kleiner et al. 2007; Pelli 1997). This computer in turn received commands from a real-time I/O system from National Instruments (Austin, USA), which ensured control of the display on a frame-by-frame basis, as well as real-time monitoring of animal behavior and reward. The system was described recently (Chen and Hafed 2013; Tian et al. 2018; 2016).

### Behavioral tasks

#### Experiment 1: Temporal frequency series

The monkeys fixated a small white spot for 350-550 ms at trial onset, after which the spot started moving either horizontally or vertically along a sinusoidal position trajectory of only 30 min arc (0.5 deg) amplitude. The monkeys had to track this moving target with their eyes, and target position in deg (along either the horizontal or vertical axis) could be described by the following equation

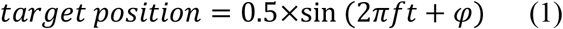

where *f* is the temporal frequency of target trajectory, *t* is time from motion onset, and *φ* could be either 0 or π across trials. The temporal frequency, *f*, of target motion was chosen randomly from trial to trial from among the following values: 0.1, 0.2, 0.3, 0.4, 0.5, 0.6, 0.8, 1, 2, 3, 4, or 5 Hz. Target motion duration was constant within a session, but could vary across sessions in the range of 3000-4200 ms, depending on animal motivation on any one day. In all cases, we had a long enough target motion duration to ensure that we were analyzing steady-state tracking behavior, even for the smallest values of *f* (associated with the slowest target position changes). Horizontal and vertical target trajectories were collected in different blocks of trials, and we analyzed a total of 867 trials from monkey A and 1392 trials from monkey M. We did not penalize the monkeys (e.g. by aborting trials) for making catch-up saccades as long as they stayed within a radius of ~1-1.5 deg around the instantaneous target position.

#### Experiment 2: Amplitude series

The monkeys performed a similar experiment to that described above, but this time, the temporal frequency, *f*, was maintained at 0.5 Hz. Also, we interleaved different target motion directions, and we varied the amplitude of the target motion. Since 0.5 Hz was sufficiently high to allow turnarounds in pursuit direction at reasonable times in a trial, we relaxed trial length in this experiment to 1700-2000 ms. Target motion trajectory was now described by the following equations

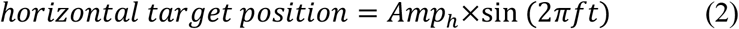

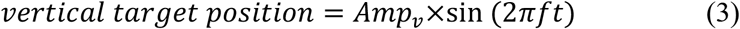

where *f* was fixed at 0.5 Hz, and *Amp_h_* and *Amp_v_* specified the target motion amplitudes. For purely horizontal target motions, *Amp_h_* and *Amp_v_* were chosen to result in radial amplitudes of 0.25, 0.5, 1, or 2 deg. We also introduced two oblique directions: “+45 deg oblique pursuit” was used to describe the case when *Amp_h_* and *Amp_v_* were equal and positive (again from among 0.25, 0.5, 1, or 2 deg); “+135 deg oblique pursuit” was used to describe the case when *Amp_v_* had an opposite sign from *Amp_h_* and *Amp_h_* was negative (e.g. −0.25 deg for *Amp_h_* and +0.25 deg for *Amp_v_).* In other words, the two oblique directions both started with upward motion trajectory, but with one first moving rightward/upward (+45 deg) and the other first moving leftward/upward (+135 deg). During the sessions, we interleaved all amplitude and direction conditions. We analyzed a total of 969 trials from monkey A and 2145 trials from monkey M.

#### Experiment 3: Fixation comparison

For a subset of analyses, we compared eye velocity during tracking of small-amplitude motion trajectories to eye velocity during fixation. The monkeys simply fixated the same small spot for 1000-1400 ms before getting rewarded. We analyzed a total of 3160 trials from monkey A and 5222 trials from monkey M.

### Data collection strategy

When still naïve, the animals were initially trained on fixation and simple visually-guided saccades. This was necessary in order to be able to calibrate them (see below), but it was also, in reality, near-automatic behavior emerging naturally from the animals themselves (i.e. requiring minimal training): they naturally fixated small spots on an otherwise gray background. The monkeys were then trained on slightly more demanding saccade tasks necessary for neurophysiology in oculomotor circuits (e.g. delayed visually-guided and memory-guided saccade tasks). The smooth pursuit tasks of the present study were introduced early on in training, and also intermixed with much faster (e.g. 12-20 deg/s speeds), constant-speed smooth pursuit along the horizontal direction (e.g. Buonocore et al. 2018). In our experience, the ocular following behavior that we observed in the present study was again near-automatic, because there was only a single spot on the display, and because the animals were aware that looking at this spot rewards them at the end of the trial. Typically, we intermixed blocks of 200-300 trials of pursuit (from either Experiment 1 or 2) and blocks of 200-300 trials of fixation (Experiment 3), often with the other tasks described above happening earlier and later in the session. The monkeys routinely performed a total of 1000-2000 trials per session (at a rate of ~1000 trials per hour), with short breaks introduced in between small blocks of 200-300 trials. Moreover, since the smooth pursuit trials of the present study could be run at different times in a session (early versus late), we did not notice a substantial factor of fatigue on performance in the data that we present here (e.g. in the proportion of rewarded trials or the distribution of reaction times in the different saccade tasks).

### Data analysis

#### Detecting catch-up saccades

We converted raw eye tracking measurements to calibrated eye rotations using the methods described in (Tian et al. 2016). Briefly, at the beginning of every experimental session, the monkeys were presented with a calibration task. The monkeys fixated a spot presented at one of 19 different locations for at least 700 ms. The locations covered screen center, six locations along the horizontal meridian relative to screen center, four locations along the vertical meridian, and 8 locations along either +45 or −45 oblique directions. Each location was sampled at least 6 times, and we then computed the best fitting parameters for polynomial functions relating calibrated eye positions to raw horizontal and vertical eye tracker measurements (Tian et al. 2016).

We then detected catch-up saccades using velocity and acceleration criteria (Chen and Hafed 2013; Hafed et al. 2009; Krauzlis and Miles 1996), and we manually inspected all movements to correct for misses or false detections. For a subset of the data, we used instead a novel state-of-the-art machine-learning approach for saccade detection using convolutional neural networks, which we have recently developed (Bellet et al. 2018).

Catch-up saccade detection was necessary for performing focused analyses on smooth pursuit eye movements (see below), but we also analyzed interesting properties of these saccades themselves. For example, we explored both the frequency of occurrence and amplitude of these eye movements as a function of either target motion temporal frequency, *f*, or amplitude (*Amp_h_, Amp_v_*). We also analyzed whether catch-up saccades were correcting for position error or retinal slip (velocity error) when they occurred. Specifically, we measured position or velocity error between the eye and target immediately before or after a given catch-up saccade during steady-state pursuit. If the error was smaller after the saccade, then the saccade was classified as being corrective (for either position or velocity error). During pursuit initiation at trial onset (e.g. Fig. 10), we also analyzed first catch-up saccade amplitude and related it to the existing position error at saccade onset (e.g. Fig. 10D).

#### Measuring smooth pursuit gain and phase lag

We plotted eye velocity as a function of time for either horizontal or vertical eye movements. For oblique motion trajectories (Experiment 2), we performed a coordinate rotation such that one component of eye velocity was along the motion trajectory and the other was orthogonal, and we plotted eye velocity for the component along the motion trajectory (in other analyses, we also analyzed the horizontal and vertical components of oblique eye velocity independently). We then picked a period of steady-state smooth pursuit execution by excluding the first 1000 ms of eye movement data after target motion onset (in Experiment 2, we relaxed this to 300 ms primarily because the trials were shorter in Experiment 2 than in Experiment 1). In each velocity trace, we then excised any interval during which a catch-up saccade was being executed, and we also removed 10 ms before and 10 ms after each such saccade. For visualization purposes (e.g. Fig. 1), the data that were excised were replaced by Not-a-Number (NaN) labels, such that averages of eye velocity across trial repetitions of a given condition did not include the large velocity transients associated with catch-up saccades. For fitting purposes (see below), we replaced the excised data by linear interpolations between saccade onset and end.

**Figure 1.**
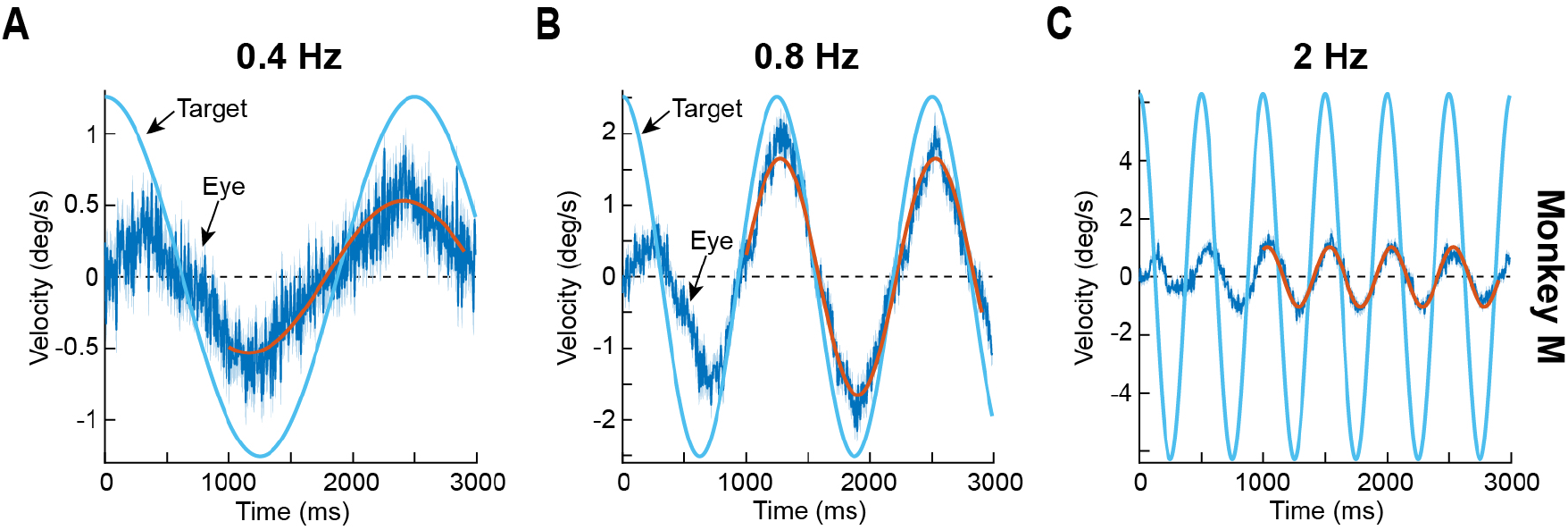
Band-pass nature of monkey smooth pursuit of small-amplitude slow motion trajectories. (**A**) Example tracking behavior from monkey M with 0.4 Hz sinusoidal target motion. The solid blue sinusoid represents target velocity. The dark blue data plot shows mean eye velocity across trials (surrounded by 95% confidence interval bands in a fainter color). The red sinusoid is a fit of the eye velocity data in a sustained interval starting 1000 ms after target motion onset. As can be seen, the eye tracked the frequency of target motion well, but with a markedly low peak velocity (i.e. low gain). (**B**) Tracking gain was significantly higher at 0.8 Hz. This panel is formatted identically to **A**. (**C**) For even higher frequencies, pursuit gain decreased again, as evidenced by the much smaller amplitude of the sinusoid describing eye velocity relative to that describing target velocity (red and blue sinusoids, respectively). Note that phase lag also increased (compare the phase of the solid red and blue sinusoids in each panel). Error bars, when visible, denote 95% confidence intervals. N = 59, 58, and 58 trials for 0.4 Hz, 0.8 Hz, and 2 Hz, respectively.

After plotting saccade-free eye velocity from any one trial, we fitted the resulting curve, using a least-squares fitting algorithm, with a sinusoidal function (of appropriate temporal frequency for the condition) in which the amplitude and phase values of the sinusoid were the fitting parameters. This resulted in a population of amplitudes and phases from the fitting procedure across trial repetitions of a given condition. For example, for 0.5 Hz horizontal target trajectories in Experiment 1, we could have a population of *N* fitted gains or phase lags across trials. We then summarized these *N* values into the mean gain or mean phase lag at 0.5 Hz target motion temporal frequency, and with appropriate 95% confidence intervals. “Gain” was defined as the ratio of the fitted eye velocity amplitude in a sinusoid divided by the true amplitude of the target velocity sinusoid. For example, for*f* Hz horizontal target trajectory with *Amp_h_* deg position amplitude in equation 1 above (and also equations 2 and 3), the target velocity amplitude was 2*πf Amp_h_*. Similar procedures were performed for all conditions. In all analyses, we had >25 trials per condition in each animal (in most cases, many more; e.g. see Fig. 2). We also excluded from our analyses the 0.1 Hz target motion frequency condition. Even though we could obtain similar goodness of fit values to other temporal frequencies when computing average sinusoids of eye velocity, the variance in the fitted sinusoids across individual trials was large enough to reduce our confidence in estimating the true effect of this slow motion frequency. The monkeys were also frequently using microsaccades to maintain gaze on the target. Inspection of the raw data, however, does not rule out making a reasonable inference that this frequency had similar effects on smooth pursuit gain to the 0. 2 Hz frequency condition, which we do present in our results (and which are consistent with Cunitz, 1970 in humans).

In a subset of analyses (e.g. Fig. 10), we measured eye velocity directly. For example, we estimated eye velocity during smooth pursuit initiation or during baseline fixation (before target motion onset). We defined a measurement interval of 50 ms, starting either at −100 ms or +100 ms from motion onset. The earlier interval measured eye velocity during fixation, whereas the latter interval measured eye velocity during smooth pursuit initiation.

#### Spectral analysis of eye velocity

For some analyses of the data from Experiment 1, we performed a discrete Fourier transform decomposition of eye velocity traces from the different values of*f* in equation 1. We picked, in each trial, an epoch of steady-state smooth pursuit (i.e. removing the initial component immediately after motion onset as described above) that was 2700 ms long, and we replaced periods during catch-up saccades with linear interpolations of eye velocity between pre- and post-saccade points. We then applied a Hanning window followed by discrete Fourier transformation to investigate whether low-gain tracking was still modulated by the temporal frequency of the target motion. We then plotted the average spectrum across the individual trial spectra.

Unless otherwise specified, error bars in the figures presented in this paper designate 95% confidence intervals, allowing the statistical robustness of our results to be easily assessed.

## Results

### Band-pass tuning for small-amplitude slow motion tracking in the monkey

Our goal was to systematically characterize the quality of monkey ocular control when tracking small-amplitude slow motion trajectories. We were motivated by the more general question of how slow ocular drifts that occur during gaze fixation may be controlled, and how similar such control may be to the control needed when volitionally tracking a moving target. We therefore asked two monkeys to pursue a small spot that moved sinusoidally with an amplitude of only 30 min arc (0.5 deg) and different temporal frequencies (Experiment 1; Material and Methods). For temporal frequencies of <1 Hz, the peak velocities of target motion (based on equation 1) were always <3.14 deg/s, and they were smaller for even lower frequencies like 0.4 Hz (1.26 deg/s), and also at off-peak-velocity epochs of tracking. Therefore, the target velocities involved in our experiments were similar in scale to the velocities with which the eye may drift on its own during steady fixation (Cherici et al. 2012; Martins et al. 1985).

We found that eye velocity always tracked the temporal frequency of the target, albeit to varying degrees of success. For example, Fig. 1A shows average saccade-free (Materials and Methods) eye velocity from monkey M when this monkey tracked a horizontally moving spot at 0.4 Hz in Experiment 1. Error bars denote 95% confidence intervals across trials, and the solid blue line shows the true target velocity (based on the derivative of equation 1). As can be seen, the eye moved sinusoidally at a similar temporal frequency to the target, but eye velocity gain was low; fitting a sinusoid to the eye velocity data (in the sustained pursuit interval; Materials and Methods; red line in Fig 1A) showed a peak velocity amplitude in the fit of 0.532 deg/s relative to the true target peak velocity of 1.26 deg/s, resulting in a gain of 0.4235. There was also a phase lead of ~90 ms, which amounted to a lead of 0.036 of a full cycle of motion trajectory (3.6% of a full cycle). When the target temporal frequency was 0.8 Hz instead, eye velocity gain was higher at 0.708 (Fig. 1B), but it then decreased once again for even higher frequencies (e.g. Fig. 1C; temporal frequency of 2 Hz; gain = 0.163). Phase lead or lag also matched the gain changes by progressively shifting towards larger and larger lags, with 0.8 Hz showing now a minimal phase delay (26 ms, or 0.021 of a full cycle) in tracking (as opposed to a lead at 0.4 Hz), and the higher frequency showing an even more substantial delay of 35 ms or 0.07 of a full cycle. Therefore, monkey smooth ocular tracking of small-amplitude slow motion trajectories may be described as being band-pass in nature, unlike earlier descriptions of smooth pursuit tuning (with much faster target speeds) as being low-pass (Collewijn and Tamminga 1984; Fabisch et al. 2009; Rottach et al. 1996). This difference is not necessarily due to the use of monkeys in our current study, as opposed to humans in the earlier ones, because monkeys are indeed capable of high-gain sinusoidal pursuit of foveal spots when faster target speeds (but similar low temporal frequencies) are used (Hafed et al. 2008; Hafed and Krauzlis 2008). It is very intriguing to us, nonetheless, that very highly trained human subjects performed significantly better at low temporal frequencies than our monkeys when faced with similar small-amplitude motions (Martins et al. 1985). Concerning phase lags, the observations above seem to be in line with earlier observations with larger-amplitude motion trajectories (Collewijn and Tamminga 1984; Rottach et al. 1996).

We confirmed the band-pass nature of small-amplitude slow motion tracking in our two monkeys, and also with both horizontal and vertical tracking. For each temporal frequency, *f*, in equation 1 (Materials and Methods), we estimated the gain of pursuit (similar to Fig. 1; Materials and Methods) and plotted it for horizontal and vertical tracking, along with 95% confidence intervals (Fig. 2A). In both monkeys, pursuit gain depended on temporal frequency (p<10^-69^, 1-way ANOVA in each monkey and for either horizontal or vertical tracking). Moreover, gain peaked near 1 Hz for both horizontal and vertical tracking (mean +/- s.d.: monkey M, peak gain was at 1 Hz; horizontal tracking, 0.708 +/- 0.165 and vertical tracking, 0.326 +/- 0.163; monkey A, peak gain was at 0.8 Hz; horizontal tracking, 0.922 +/- 0.183 and vertical tracking 0.429 +/- 0.183). Vertical tracking had significantly worse pursuit gain than horizontal tracking (Rottach et al. 1996) for all target motion frequencies between 0.3 Hz and 1 Hz in monkey M, and up to 3 Hz in monkey A, as confirmed by t-tests (p<0.05) using Bonferroni correction.

We similarly analyzed pursuit phase lag, reporting it as a fraction of a full cycle (Fig. 2B); that is, a constant temporal delay in phase would mean a larger fraction of a cycle with increasing frequencies. This was the case in both monkeys, particularly during horizontal tracking: higher target motion frequencies resulted in progressively more and more pursuit lag when represented as a fraction of a full cycle. Note that in monkey A during vertical tracking, higher frequencies were associated with an apparent phase lead (see inset in Fig. 2B at 2 Hz for monkey A), but during steady-state sinusoidal behavior, this is equivalent to a large phase lag. The insets in Fig. 2A show the total numbers of trials analyzed for each condition.

**Figure 2.**
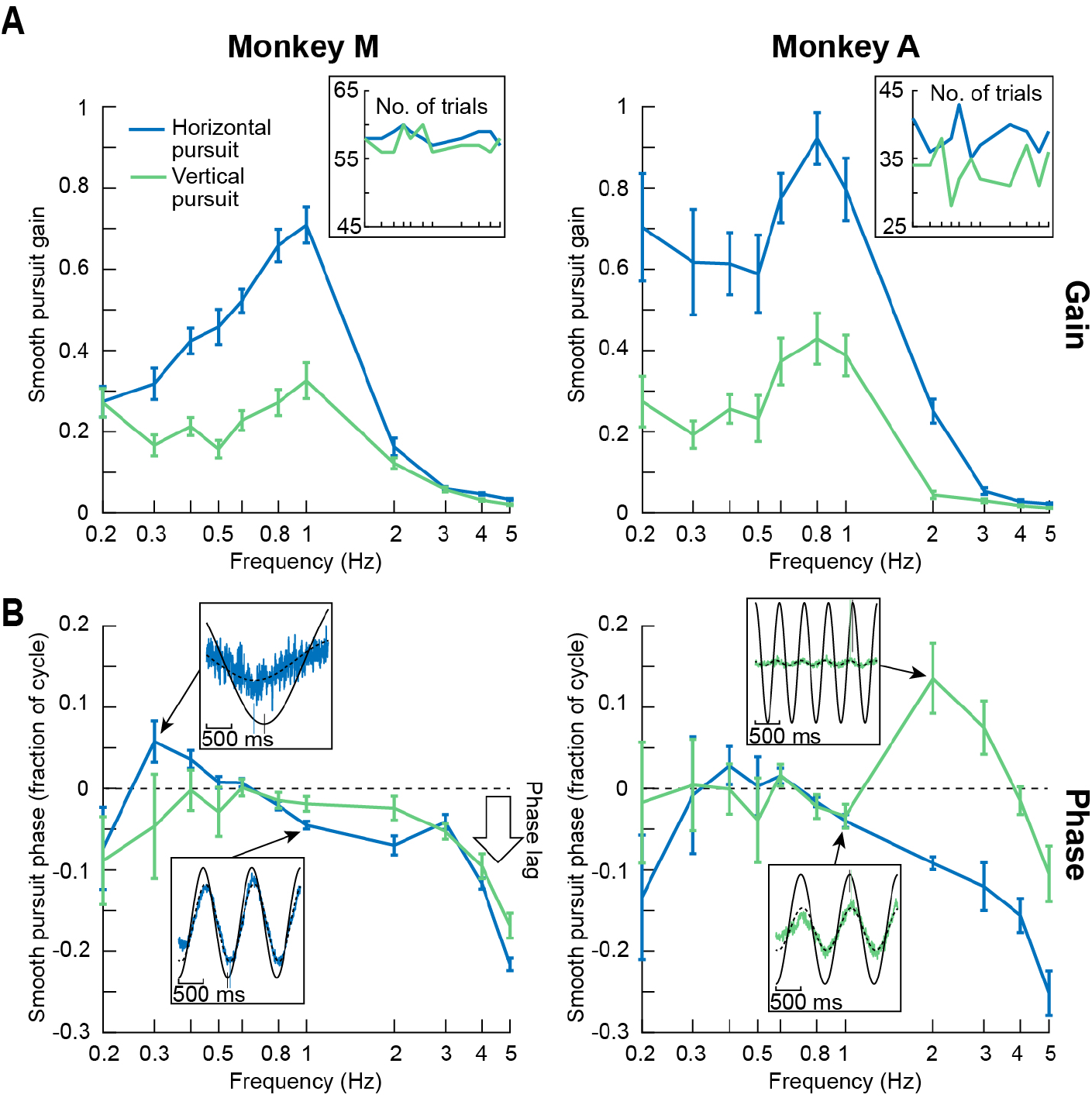
Band-pass nature of slow motion smooth pursuit in both monkeys. (**A**) Each panel shows the gain of smooth pursuit (i.e. the ratio of red sinusoid amplitude to blue sinusoid amplitude in Fig. 1) as a function of target motion frequency. The left panel shows data from monkey M, and the right panel shows data from monkey A. Error bars denote 95% confidence intervals, and the insets indicate the numbers of trials per condition used for analysis. Pursuit gain peaked at around 0.8-1 Hz for both monkeys, and was lower for both lower and higher frequencies. In monkey A, pursuit gain was slightly elevated at 0.2 Hz compared to 0.3-0.6 Hz (also true for vertical tracking in monkey M); at this frequency, the target was moving very slowly and ocular drift velocities could be substantially higher than target velocities (Martins et al. 1985; also see Cunitz 1970 and Fig. 3A). Also, in both animals, pursuit gain was significantly worse for vertical tracking as opposed to horizontal tracking. (**B**) A similar analysis but now for pursuit phase (e.g. the phase difference between the red and blue sinusoids of Fig. 1). Insets show examples of eye velocity phase relationships to target velocity curves at several representative target motion frequencies (small black and colored vertical lines clarify the phase relationship in each inset). Higher frequencies were associated with larger phase lags. Note that the phase lag is displayed here as a fraction of a cycle. Thus, a constant temporal lag would translate into a larger fraction with increasing frequency (see text). Also, note that, in steady-state sinusoidal behavior, a large phase lag could be equally interpreted as a phase lead (e.g. the inset in monkey A’s panel at 2 Hz). All error bars indicate 95% confidence intervals.

Therefore, for both horizontal and vertical smooth pursuit eye movements, tracking of very slow motion trajectories in monkeys is seemingly different from experiments with faster target speeds; for the small-amplitude slow motions, pursuit efficacy at low frequencies is significantly impaired, and only recovers at ~0.8-1 Hz. For larger target speeds, evidence from the literature shows that smooth pursuit typically exhibits low-pass behavior, with high gain at all low frequencies up to ~1 Hz (Collewijn and Tamminga 1984; Fabisch et al. 2009; Hafed et al. 2008; Hafed and Krauzlis 2008; Martins et al. 1985; Rottach et al. 1996). As stated above, in highly trained humans, such a low-pass behavior of smooth pursuit seemed to also still persist for small-amplitude target trajectories like the ones that we used (Martins et al. 1985); perhaps this difference from our monkey results is due to extensive training of the humans to avoid making catch-up saccades.

### Controllability of monkey ocular velocities as slow as those during fixational ocular drifts

Despite the relatively low gain of smooth pursuit at low temporal frequencies in Figs. 1, 2, eye velocity in our monkeys was still clearly modulated. For example, sinusoidal tracking was still evident at 0.4 Hz even with the reduced gain (Fig. 1A). As stated earlier, we were interested in this phenomenon particularly because the velocities with which tracking occurred at these low temporal frequencies was similar to the velocities with which ocular drifts during fixation normally take place. For example, peak target velocity at 0.2 and 0.3 Hz was 0.628 and 0.942 deg/s, respectively, which are all within the range of eye velocity during fixational ocular drifts (Cherici et al. 2012; Martins et al. 1985). To confirm this, we collected control fixation data from the same animals (Experiment 3). In these trials, the spot never moved, and the monkeys simply fixated it for approximately 1000 ms (Materials and Methods). We measured eye velocity during microsaccade-free fixation epochs, and we related them to the peak eye velocity at each temporal frequency during tracking (Fig. 3A). Specifically, in Fig. 3A, we plotted the same data as in Fig. 2A but now as raw measurements of peak eye velocity instead of as gain values (error bars denote 95% confidence intervals). We then plotted the average velocities observed during gaze fixation (horizontal lines), again along with 95% confidence intervals (mean +/- s.d. in deg/s: monkey M, fixational horizontal eye velocity = 0.74 +/- 0.126, fixational vertical eye velocity = 0.631 +/- 0.115; monkey A, fixational horizontal eye velocity = 0.798 +/- 0.126, fixational vertical eye velocity = 0.597 +/- 0.11). At the lowest and highest tracking frequencies (e.g. 0.2-0.3 Hz or 4-5 Hz), peak eye velocity during pursuit of slow motion trajectories was lower than eye velocity during fixation (e.g. monkey A: peak horizontal eye velocity at 0.2 Hz was 0.442 +/- 0.260 s.d. deg/s; error bars in Fig. 3A denote 95% confidence intervals). However, the eye was still well-controlled because gain was not zero (Fig. 2A). We also further confirmed this by analyzing the frequency spectrum of de-saccaded eye velocity traces for different temporal frequencies. We found that there was a peak in the power spectrum of eye velocity traces at the frequency with which the target was moving (Fig. 3B). This means that eye velocity had a harmonic component at the frequency of the target motion trajectory even at the low gain values observed in Fig. 2A. Finally, we also inspected example eye position traces for low (Fig. 3C) or high (Fig. 3D) tracking frequencies, and clear modulation of saccade-free eye position was still present (e.g. see the gray arrows in Fig. 3C, D). Therefore, slow ocular movements at velocities similar to or lower than the velocities of slow fixational ocular drifts are relatively well-controlled in this behavior (Wyatt and Pola 1981).

**Figure 3.**
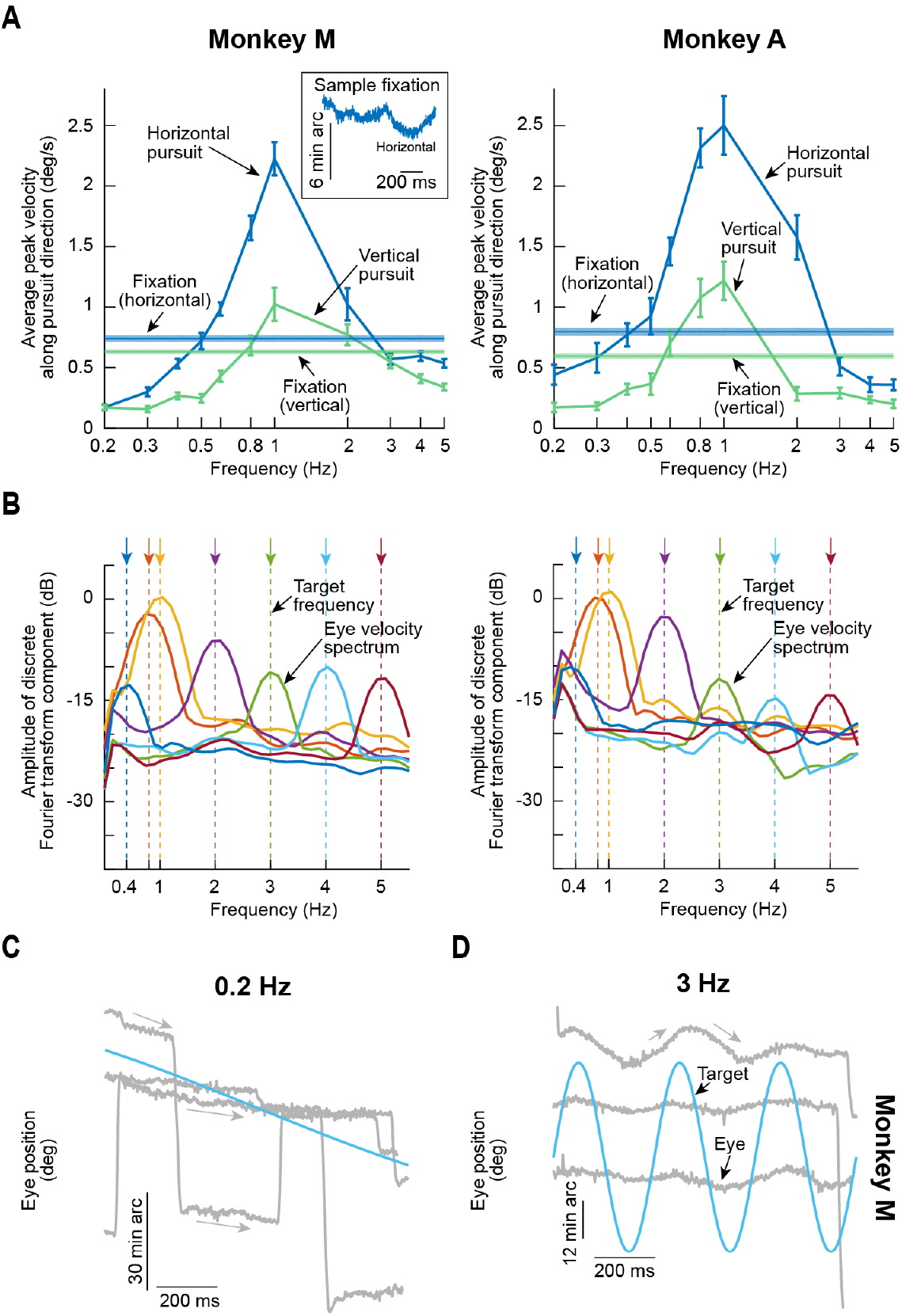
Monkey ocular velocities as slow as those during fixational drifts occurred during smooth pursuit of slow motion, but they were still systematically controlled to track the motion trajectory. (**A**) The curves show the same data as in Fig. 2A, but this time as real measurements of peak eye velocity as opposed to a gain ratio. Error bars denote 95% confidence intervals. The solid horizontal lines show average eye velocity during fixation (Materials and Methods), surrounded by 95% confidence intervals (the inset shows an example 1-second fixational eye position trace, demonstrating how ocular drift has substantial non-zero eye velocity even with a stationary fixation spot). Fixational drift velocity was higher than pursuit peak velocities at pursuit frequencies of, say, 0.2 Hz, 0.3 Hz, and 4 Hz. This means that eye velocities as slow as those during ocular drifts are controllable by the central nervous system of the monkey. (**B**) This idea is supported by analyzing the spectral content of de-saccaded eye velocity traces for different pursuit target frequencies. Even at low frequencies (e.g. 0.4 Hz and 0.8 Hz) associated with low pursuit velocities (as in **A**), eye velocity still exhibited a peak in the spectrum at a frequency near target motion. This indicates that eye velocity was still following target motion (even if at low gain). (**C**, **D**) Sample 1-second eye position traces (shown in gray) from three example trials at 0.2 Hz (**C**) or 3 Hz (**D**), both of which exhibited very weak pursuit gain (**A** and Fig. 2). In both cases, eye velocity in between saccades was systematically controlled to follow target trajectory (e.g. see the gray arrows). Thus, monkeys are capable of slow control of their drift velocities in our task.

### Dependence of monkey catch-up saccade frequency and amplitude on temporal frequency

Our results so far have focused on smooth eye velocity effects. However, we also analyzed and catalogued catch-up saccade frequency and amplitude. We found that catch-up saccades behaved in different ways for temporal frequencies higher or lower than the frequency associated with peak smooth pursuit gain (~0.8-1 Hz). For example, in Fig. 4A, we plotted the frequency of catch-up saccades as a function of temporal frequency in Experiment 1 (the faint-colored curves show the smooth velocity gain curves of Fig. 2, presented on arbitrary y-axes, to provide a reference for comparison). In both animals, catch-up saccade frequency reached a peak near the temporal frequency for which smooth pursuit gain was maximum (mean +/- s.d. in saccades/s: monkey M, at 1 Hz, horizontal trials = 2 +/- 0.466, vertical trials = 2.01 +/- 0.636; monkey A, at 1 Hz, horizontal trials = 2.79 +/- 0.631, vertical trials = 3.06 +/- 0.546). This suggests that eye position was continuously adjusted with both smooth pursuit and saccadic eye movements when overall tracking was particularly effective (i.e. with high gain). Catch-up saccade frequency then dropped for higher temporal frequencies. For example, there were only 1.19 and 1.35 saccades/s during 3 Hz horizontal tracking for monkeys M and A, respectively. This drop was not so dramatic for lower temporal frequencies, with mean saccade frequencies staying above 1.7 saccades/s in all cases up to 2 Hz target motion frequency.

These observations suggest that for the lower temporal frequencies, when the spot was barely moving, catch-up saccades played a role similar to that of fixational microsaccades: they optimized eye position on the target on average (Guerrasio et al. 2010; Ko et al. 2010; Tian et al. 2018; 2016). On the other hand, for very rapid oscillations (high frequencies), the oculomotor system was unable to keep track of the frequent flips in target position (even with saccades as opposed to smooth pursuit), and saccades were therefore more or less random events. Consistent with this, catch-up saccade amplitudes (Fig. 4B) were always small for all frequencies <1 Hz; on the other hand, catch-up saccade amplitudes increased for higher frequencies, again likely reflecting the higher position errors associated with slow smooth velocity gain.

**Figure 4.**
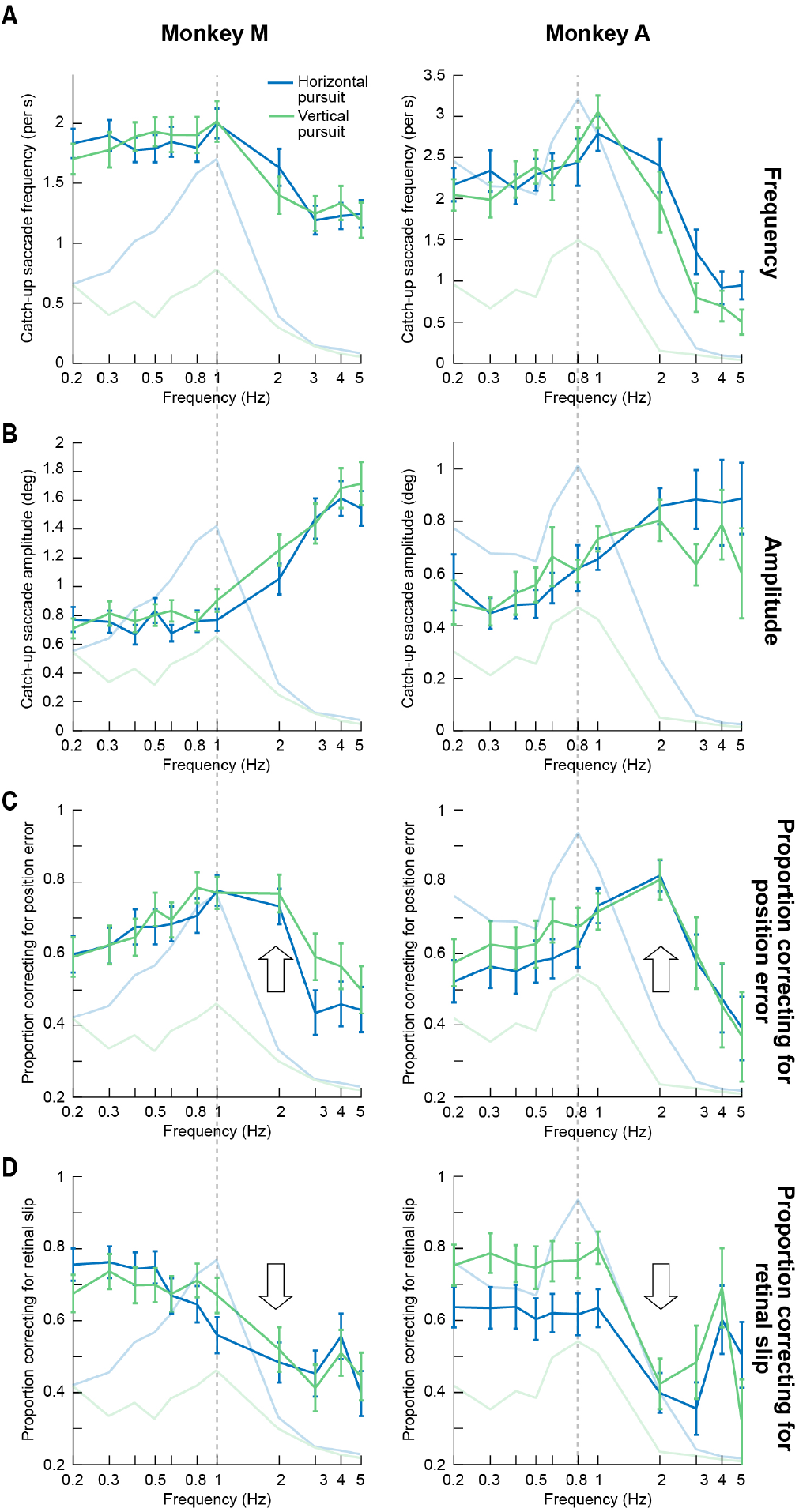
Catch-up saccades increased the effective bandwidth of ocular tracking of small-amplitude slow motion trajectories. (**A**) For both animals in Experiment 1, we estimated the frequency of catch-up saccades during steady-state smooth pursuit. Error bars (in this and all other panels) denote 95% confidence intervals, and the different colors denote horizontal and vertical tracking. The faint curves show smooth velocity gain curves from Fig. 2 (with arbitrary y-axis scaling), in the same experiment, to allow comparing catch-up saccade curves to smooth gain curves (faint dashed vertical lines denote the temporal frequencies associated with maximal smooth velocity gain in each monkey). The highest rate of catch-up saccades occurred at frequencies near those associated with maximal smooth velocity gain, with a subsequent fall-off coming at higher frequencies. The fall-off in catch-up saccade rate with increasing target motion frequency was more gradual than the fall-off in smooth velocity gain. (**B**) Catch-up saccade amplitudes increased with increasing target motion frequency, particularly during the high-frequency fall-off phase, and these patterns of results (**A** and **B**) were identical for horizontal and vertical pursuit, despite the quantitative difference in smooth velocity gain for these different pursuit directions (Figs. 2-3). Note that the vertical scales in **A** and **B** are different across the two monkeys because of their different catch-up saccade frequencies and amplitudes. (**C**) We calculated the proportion of catch-up saccades that reduced instantaneous eye position error when they occurred (Materials and Methods). In both monkeys, the likelihood of a position-error corrective catch-up saccade peaked near 2 Hz, when smooth velocity gain was already strongly reduced (upward arrows). Thus, the bandwidth of overall tracking behavior (combined smooth and saccadic eye movements) was higher than the bandwidth of smooth velocity gain alone. (**D**) A similar analysis for catch-up saccades acting to reduce instantaneous retinal slip, or velocity error, (Materials and Methods) revealed that at high frequencies (e.g. 2 Hz), catch-up saccades were not effective in reducing instantaneous retinal slip (downward arrows).

We further explored these observations by analyzing whether catch-up saccades corrected for position errors or retinal slips (i.e. velocity errors) when they occurred or not. In other words, we investigated the synergistic interactions between saccades and smooth eye movements when tracking small-amplitude slow motion trajectories. We classified each catch-up saccade as being either error-correcting or error-increasing by measuring position error or retinal slip (i.e. velocity error) after the saccade relative to before it. We found that catch-up saccades were corrective for position error even at 2 Hz when smooth pursuit gain was dramatically reduced (Fig. 4C). This indicates that catch-up saccades acted to increase the effective bandwidth of overall tracking behavior (Collewijn and Tamminga 1984); smooth velocity gain was weak but saccades corrected for position error, helping to keep the eye close to the target. In terms of retinal slip, catch-up saccades improved retinal slip when they occurred only at low frequencies (Fig. 4D). Thus, catch-up saccades served complementary roles in tracking behavior at low and high temporal frequencies.

Such complementary roles became even more obvious when we analyzed average position error and retinal slip before and after catch-up saccades for the different temporal frequencies. For example, in Fig. 5A, we plotted position error before (solid curves) and after (dashed curves) catch-up saccades during horizontal tracking. The saccades reduced position error for most frequencies <3 Hz, but they were most effective when smooth velocity gain decreased from its peak value (e.g. see the difference between dashed and solid curves at 2 Hz). This means that catch-up saccades acted to approximately equalize position error during tracking up to 2 Hz (approximately flat dashed curves). Once again, this was a higher bandwidth than the bandwidth of smooth velocity gain alone (Collewijn and Tamminga 1984). These observations were also true during vertical tracking (Fig. 5B). Conversely, catch-up saccades were most effective in reducing retinal slip only at low temporal frequencies (Fig. 5C, D).

**Figure 5.**
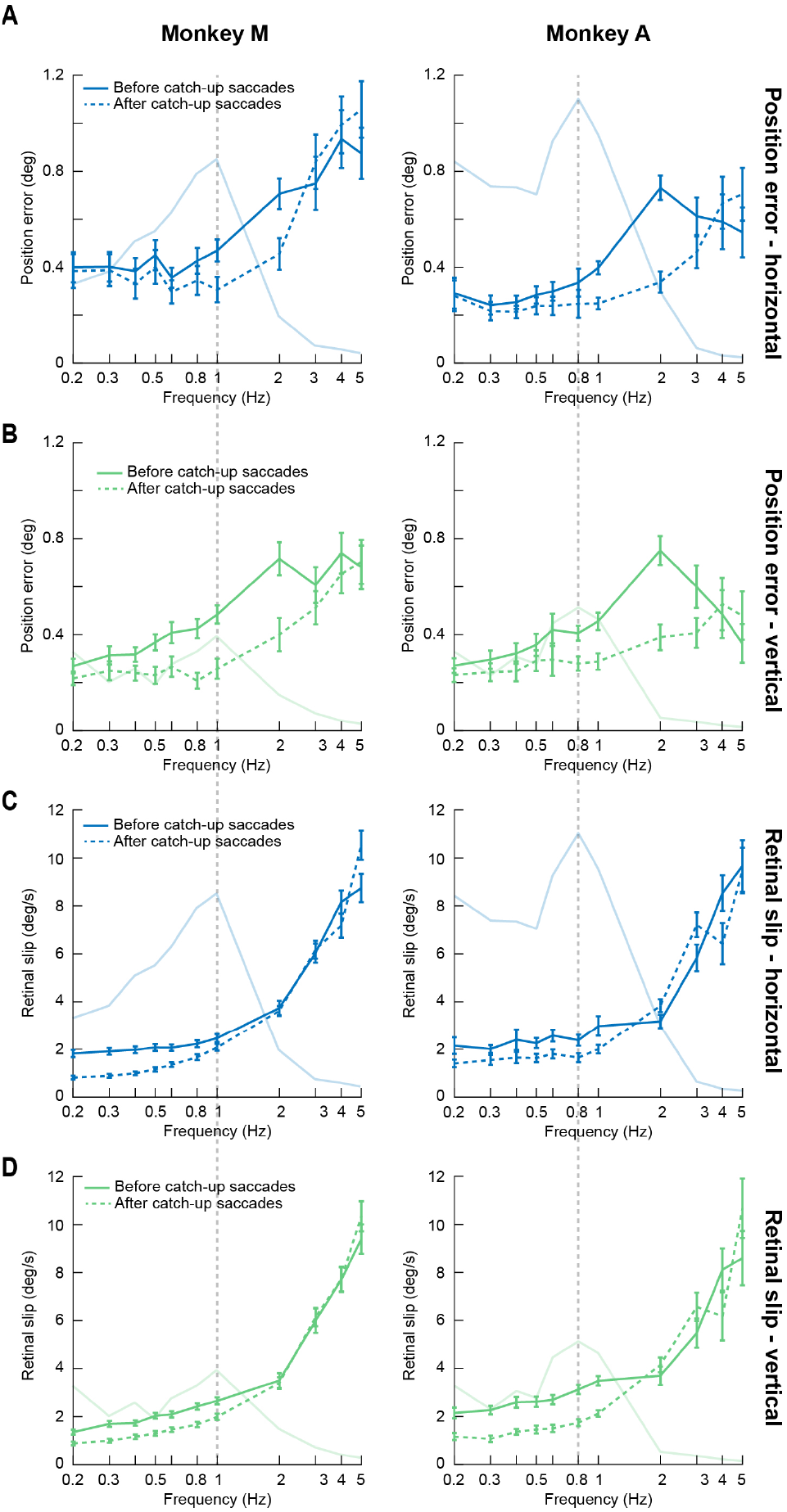
At low target motion frequencies, catch-up saccades primarily reduced instantaneous retinal slip; at high target motion frequencies, catch-up saccades primarily reduced instantaneous position error. (**A**) For each catch-up saccade in Experiment 1, we measured instantaneous position error before (solid) and after (dashed) the saccade (Materials and Methods). We then plotted these measurements as a function of target motion temporal frequency. Error bars (in this and all other panels) denote 95% confidence intervals, and the faint curves show smooth velocity gain curves for reference, exactly as described in Fig. 4. This panel shows position-error correction for horizontal tracking. As can be seen, catch-up saccades were most effective in reducing position error at temporal frequencies larger than the frequency eliciting maximal smooth velocity gain. The net effect of catch-up saccades was to equalize eye position error after catch-up saccades up to 2 Hz, even though smooth velocity gain might have been low at this high frequency. In other words, the effective bandwidth of combined smooth and saccadic tracking was higher than that with smooth velocity alone. (**B**) A similar observation was made during vertical tracking. Catch-up saccades reduced position error most effectively when smooth velocity gain started decreasing from its peak. (**C**, **D**) On the other hand, measurements of instantaneous retinal slip before and after catchup saccades revealed that catch-up saccades were most effective at reducing instantaneous retinal slip when target motion trajectories had low frequency (up to the frequency eliciting maximal smooth velocity gain). Therefore, small catch-up saccades during slow motion tracking served complementary roles at different frequencies. For target motion frequencies larger than 2 Hz, the saccades neither reduced position error nor reduced retinal slip.

All of the above interpretations are also supported by inspecting sample eye position traces from four example temporal frequencies from Experiment 1 (Fig. 6). Saccades were frequent at low temporal frequencies (Fig. 6A), and they kept the eye hovering around target location, consistent with the role of fixational microsaccades in continuously optimizing eye position (Guerrasio et al. 2010; Ko et al. 2010; Tian et al. 2018; 2016). However, a substantial fraction of them took the eye momentarily away from the target before a corrective movement was triggered. Moreover, after the saccades, retinal slip relative to target motion was reduced (e.g. gray arrows in Fig. 6A following a trajectory similar to the target direction in the blue sinusoid). On the other hand, at an optimal frequency for smooth velocity gain (Fig. 6B), position error was small most of the time due to the fact that smooth velocity gain was high, as well as the fact that the slightly more frequent catch-up saccades (relative to Fig. 6A) were primarily corrective movements. When frequency increased further to 2 Hz (Fig. 6C), smooth velocity gain was poorer, but catch-up saccades were frequent and kept the eye, on average, hovering near the target. Thus, the saccades compensated for the smooth velocity loss and increased the effective bandwidth of tracking in terms of position error. Finally, saccades at even higher temporal frequencies (Fig. 6D) were less frequent, large, and very often increasing eye position error, rather than decreasing it, and for substantial amounts of time (>500 ms).

It is also interesting to note that despite the large difference in smooth pursuit gain between horizontal and vertical tracking in Experiment 1 (Fig. 2), catch-up saccade frequency and amplitude were not that different from each other across tracking directions (Fig. 4). This might suggest that there is a larger tolerance for oculomotor errors along the vertical dimension, perhaps because of potential asymmetries in oculomotor circuits (Hafed and Chen 2016), although this remains to be just a hypothesis at the moment.

**Figure 6.**
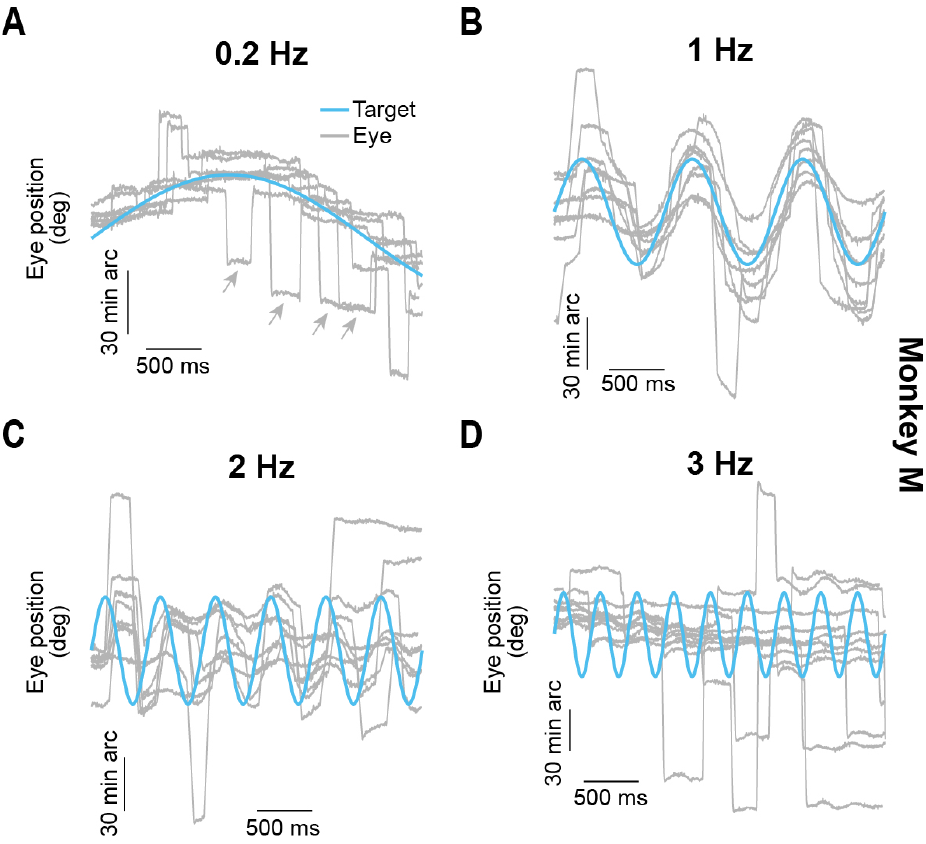
Monkey catch-up saccades at low, intermediate, and high temporal frequencies in Experiment 1. (**A**) Example eye position traces (gray) when tracking a target moving at 0.2 Hz (true target position is shown by the blue sinusoid). Smooth velocity (i.e. slow ocular drifts) tracked target motion, albeit at a relatively low gain (Fig. 2), and there were plenty of catch-up saccades. Thus, the slow eye movements here appeared similar to fixational ocular drifts in terms of velocity (e.g. inset in Fig. 3A). Moreover, catch-up saccades were frequent, and resulted in the eye “hovering” around target position on average, as with fixational microsaccades. Even though some saccades increased eye position error, retinal slip after them was aligned with target speed (e.g. gray arrows; also see Fig. 5). (**B**) Example traces in the same format as that in A, but for 1 Hz target motion frequency. Smooth velocity gain was high (Fig. 2), and also catch-up saccades were effective in correcting for eye position errors in most cases (Figs. 4, 5). (**C**) By 2 Hz, the smooth velocity gain was low again (Fig. 2), but the eye was still well-localized near the target most of the time, and this was due to the frequent position-error correcting catch-up saccades (Figs. 4, 5). Thus, 2 Hz was still within the effective bandwidth of combined saccadic and smooth tracking (Figs. 4, 5). (**D**) At even higher frequencies, slow movements also tracked the target motion (at very low gain; Fig. 2), but this time, catch-up saccades were less frequent (Fig. 4), and they were large, often deviating the eye substantially away from the target, and for substantial periods of time (>500 ms) (Fig. 5).

### Over-tracking of the horizontal component of oblique small-amplitude slow motion trajectories relative to the vertical component

We also sought to compare the effects of temporal frequency that we observed above to those of movement amplitude and direction for a given frequency. We therefore conducted Experiment 2 (Materials and Methods) in which temporal frequency was pegged at 0.5 Hz but movement amplitude varied between ~15 min arc (0.25 deg) and ~2 deg (Materials and Methods). Movement direction also included oblique tracking (Materials and Methods). Even though 0.5 Hz was a slightly sub-optimal temporal frequency in terms of smooth pursuit gain (e.g. Fig. 2), we chose it to maintain the slowest possible motion trajectories throughout our experiments. Also, our previous results (e.g. Fig. 3) demonstrated that tracking was still possible at this frequency. Overall, we found expected results in terms of smooth pursuit gain as a function of target motion trajectory amplitude. For example, for both cardinal (horizontal and vertical) and oblique pursuit, smooth pursuit gain increased with increasing target motion amplitude (Fig. 7). This might explain why our results from Experiment 1 above showed band-pass behavior (Fig. 2) when the target amplitude was small; this band-pass effect was primarily due to the very small motion trajectory amplitudes (and, correspondingly, velocities) used when compared to studies with large-amplitude pursuit, and increasing the target motion amplitude in the current experiment has alleviated this.

**Figure 7.**
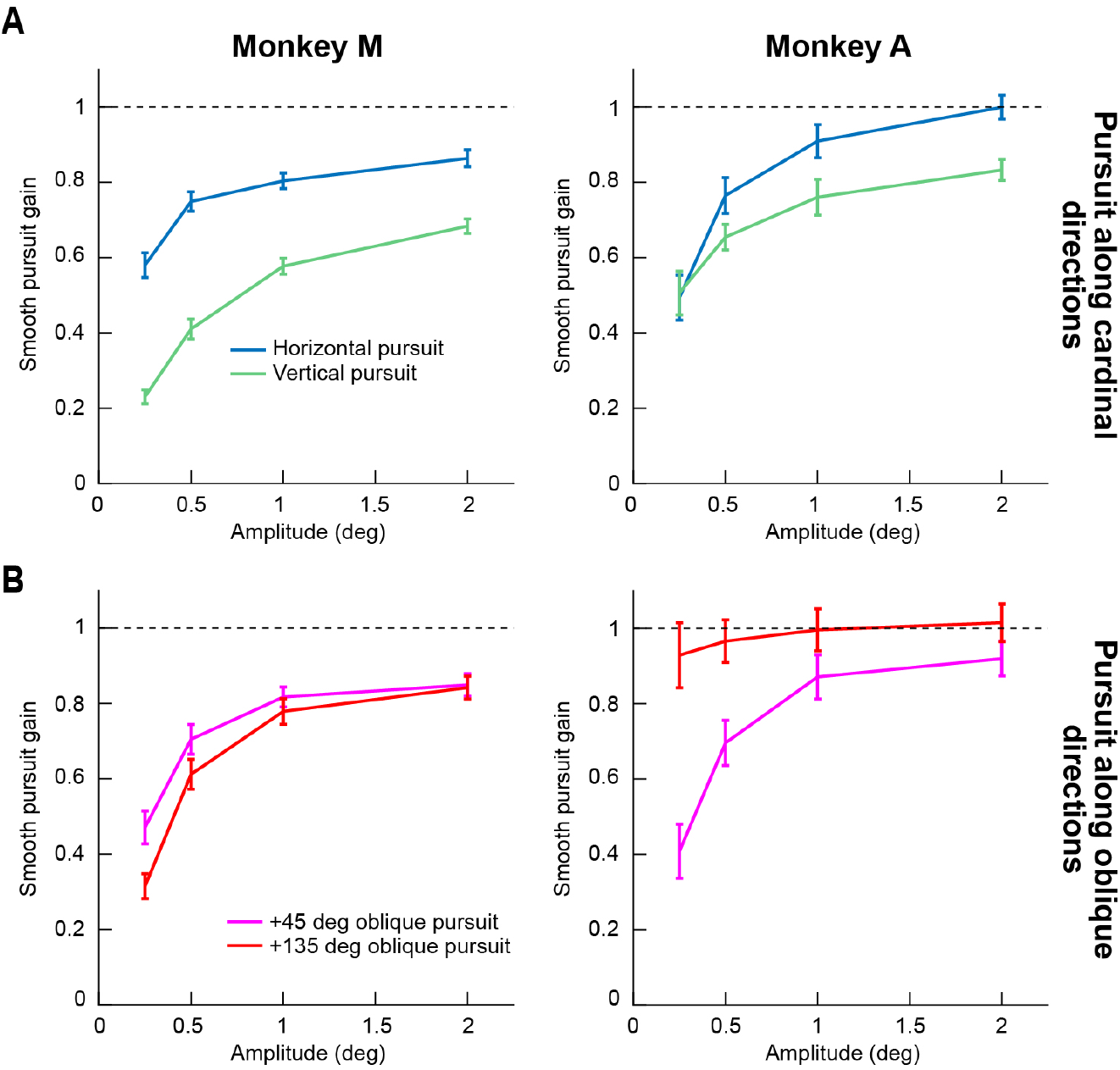
Higher pursuit gain for tracking larger and larger movement amplitudes. (**A**) In Experiment 2, we varied target sinusoidal position amplitude, and also introduced oblique pursuit trajectories. For cardinal directions, we obtained expected results based on what we found in Experiment 1. Pursuit gain was higher for horizontal than vertical pursuit, and pursuit gain progressively increased with increasing target amplitude for both horizontal and vertical pursuit. Note that gain seemed to asymptote near a value of 1. (**B**) For oblique pursuit directions, there was still an increase in pursuit gain with target amplitude. Note that we labeled target amplitude on the x-axis in this panel by the amplitude of either the horizontal or vertical component. In reality, the real radial amplitude was slightly larger because of the combined horizontal and vertical amplitudes (Materials and Methods), but we used similar labels to **A**, just in order to facilitate readability across the panels. The basic effect of increasing gain with increasing target motion amplitude is the same. All error bars denote 95% confidence intervals.

We also analyzed the horizontal and vertical components of oblique pursuit independently. We found that the horizontal component consistently had higher gain than the vertical component (Fig. 8). These results are similar to observations with larger-amplitude pursuit in humans (Rottach et al. 1996). These results, for the horizontal component at least, are also reminiscent of overshoot in visually-guided saccade amplitudes in humans for very small retinal eccentricities (Kalesnykas and Hallett 1994).

**Figure 8.**
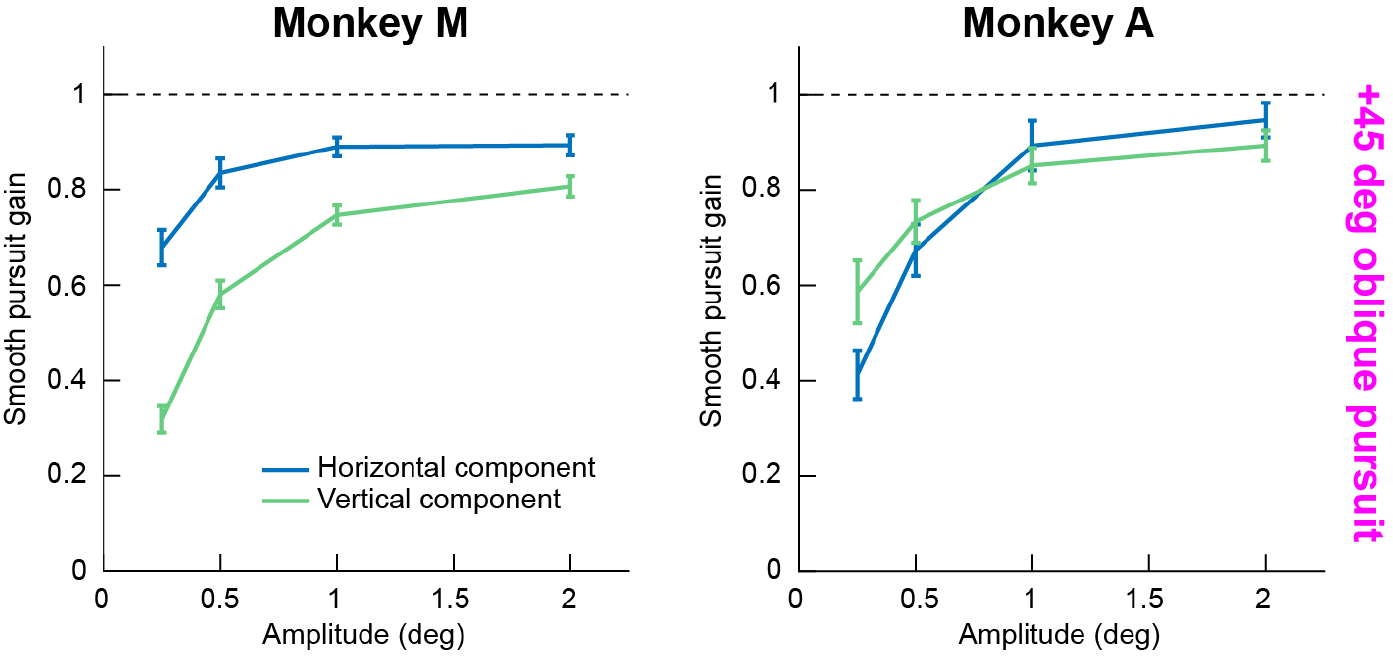
Over-tracking in the horizontal dimension, relative to the vertical dimension, during smooth pursuit of small-amplitude oblique motion trajectories. We investigated the oblique pursuit conditions in Fig. 7B more closely by plotting the horizontal and vertical components of eye velocity separately for an example oblique pursuit direction. In both animals, pursuit gain was higher in the horizontal component of eye velocity than the vertical component. Once again, the labels on the x-axis denote the amplitude of either the horizontal or vertical component for simplicity (see the legend of Fig. 7B above for reasons why). All error bars denote 95% confidence intervals.

The oblique motion trajectories in Experiment 2 were also accompanied by larger catch-up saccades for these trajectories. In Fig. 9, we plotted catch-up saccade frequency (Fig. 9A) and amplitude (Fig. 9B), as we had done earlier for Experiment 1. Consistent with Experiment 1, increased smooth pursuit gain was associated with an increase in catch-up saccade frequency (Fig. 9A), suggesting synergistic interactions between smooth pursuit eye movements and saccades to optimize eye position on the target (de Brouwer et al. 2002); this is similar to us seeing the most catch-up saccades in Experiment 1 for the temporal frequencies (0.8-1 Hz) in which smooth velocity gain was also at a maximum. In terms of catch-up saccade amplitude, it also increased with increasing target motion amplitude; in addition, while an increase in saccade amplitude was expected with increasing target position trajectory amplitude, the increase was stronger for oblique directions (Fig. 9B). For example, in monkey A, both oblique directions had higher saccade amplitudes than horizontal or vertical trajectories for 2 deg target motion amplitudes (mean +/- s.d.: +45 deg tracking, 1.14 +/- 0.601; +135 deg tracking, 1.08 +/- 0.641; vertical tracking, 0.879 +/- 0.453; horizontal tracking, 0.804 +/- 0.381), and in monkey M, one of the oblique directions did (+45 deg tracking, 1.08 +/- 0.531; vertical tracking, 0.916 +/- 0.407; horizontal tracking, 0.817 +/- 0.451). This result might reflect the slightly higher peak velocities associated with oblique tracking in our task design (for example, oblique tracking with a horizontal and vertical amplitude of 2 deg each meant an overall radial amplitude of 2.82 deg; Materials and Methods).

**Figure 9.**
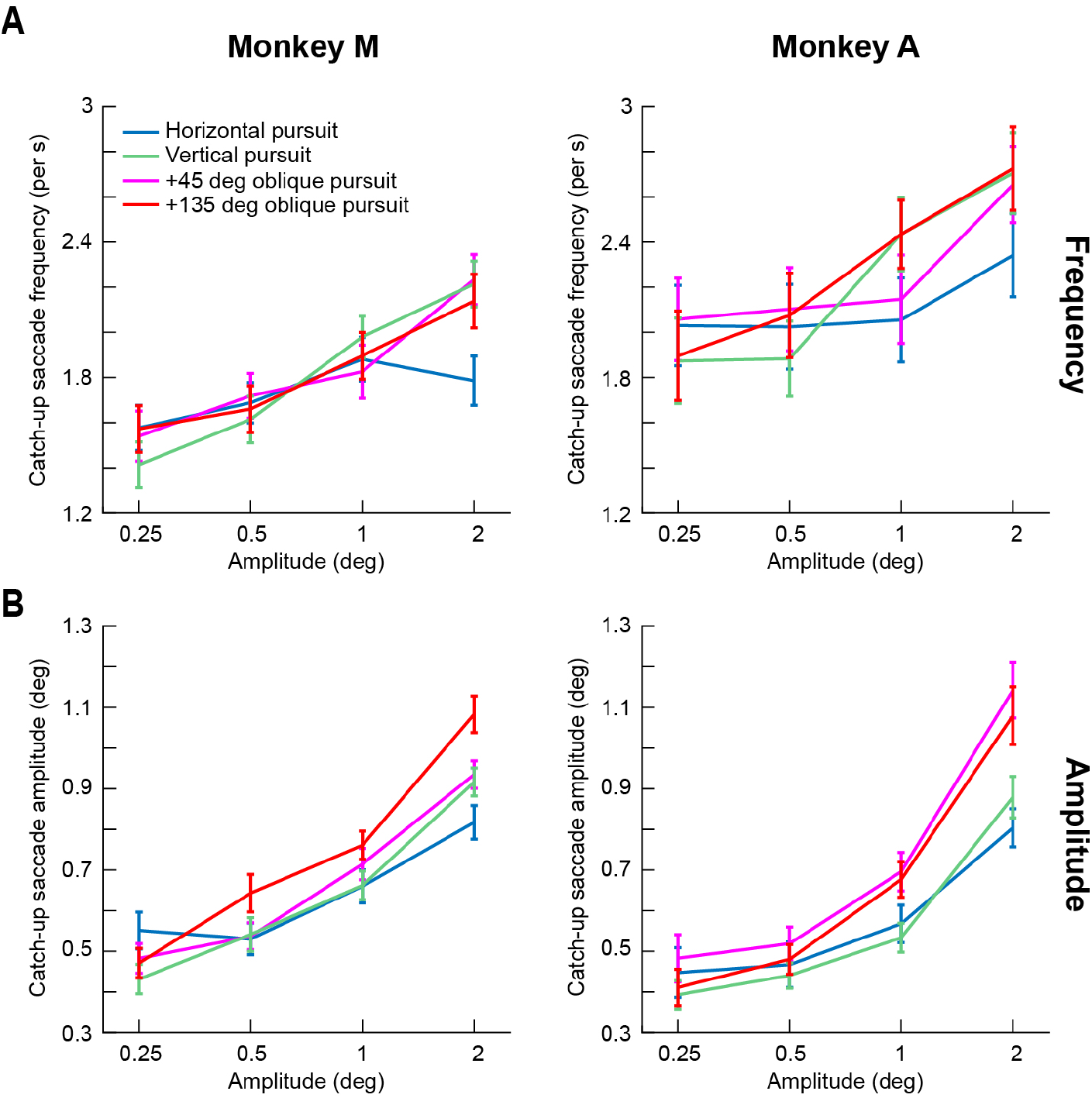
Catch-up saccade frequency and amplitude in Experiment 2. (**A**) Consistent with Experiment 1, when smooth velocity gain was high (Figs. 7-8), catch-up saccade frequency was high. (**B**) Catch-up saccade amplitude also expectedly increased with larger target motion excursions. All error bars denote 95% confidence intervals.

### Dependence of monkey first catch-up saccade latency and smooth pursuit initiation on target motion

We also analyzed the properties of the very first catch-up saccade during smooth pursuit initiation, as well as the smooth component of initial eye acceleration itself. For all saccades (including fixational microsaccades) occurring in the interval 0-300 ms after target motion onset, we plotted these movements’ amplitudes as a function of their occurrence time. We also plotted either baseline eye velocity (in the 50 ms interval starting at −100 ms from target motion onset) or smooth pursuit initiation eye velocity (in the 50 ms starting 100 ms after target motion onset; ensuring no saccades within each interval). We observed expected relationships between initial catch-up saccades and initial smooth pursuit eye velocity. For example, in Experiment 2, with 0.5 Hz target trajectory variation, eye position error of the target (relative to initial fixation location if the eye did not start tracking) monotonically increased in the first 300 ms of any trial (and up to 500 ms). Therefore, if a saccade were to occur during initiation (and there was no associated smooth acceleration after motion onset), then saccade amplitude was expected to increase with increasing time after motion onset (since the position error at saccade triggering would be larger). However, this was not always obvious in the data for the smallest target amplitudes (Fig. 10A; only horizontal tracking data is shown for clarity). We think that this is so because of the small amplitude of position errors associated with the initial target motions in this experiment, and also because of a concomitant increase in smooth eye velocity to track the target (Fig. 10B; only horizontal tracking data is shown for clarity). In other words, after motion onset, the eye often started to accelerate smoothly, therefore already reducing eye position error. Such reduction may have alleviated the need to increase first catch-up saccade amplitude. Only when target position amplitude was large enough (2 deg) did there arise a need for increasing initial catch-up saccade amplitude (peak amplitude, mean +/- s.d. deg: monkey M, occurring 270-300 ms from trial onset, 1.4 +/- 0.247 deg; monkey A, 120-150 ms from trial onset, 1.53 +/- 0.3 deg). For such a larger position amplitude of the motion trajectory, even the initial component of smooth pursuit acceleration was not sufficient to reduce eye position error sufficiently; a larger saccade was therefore necessary. This idea is illustrated in Fig. 10C showing raw pursuit velocities with saccades excised from the averages (i.e. replaced by Not-a-Number labels) as per Materials and Methods (again shown from horizontal tracking only for clarity). Pursuit initiation velocity increased for trajectories with amplitudes of 2 deg compared to, say, 0.5 deg. However, the increase in velocity did not necessarily allow for completely eliminating eye position error, resulting in the need for an initial catch-up saccade whose amplitude gradually increased with increasing time after motion onset (Fig. 10A; 2 deg motion position amplitude). Therefore, there was a synergistic interaction between smooth pursuit initiation and initial catch-up saccade execution, which also likely occurs between slow ocular drifts and microsaccades (Chen and Hafed 2013). Note that in Fig. 10A, saccades occurring <50-60 ms after target motion onset were fixational microsaccades and not really target-driven catch-up saccades because they occurred too early to reflect the new visual error signal introduced by target motion onset. This is why these movements were also small in amplitude even for 2 deg motion amplitudes.

**Figure 10.**
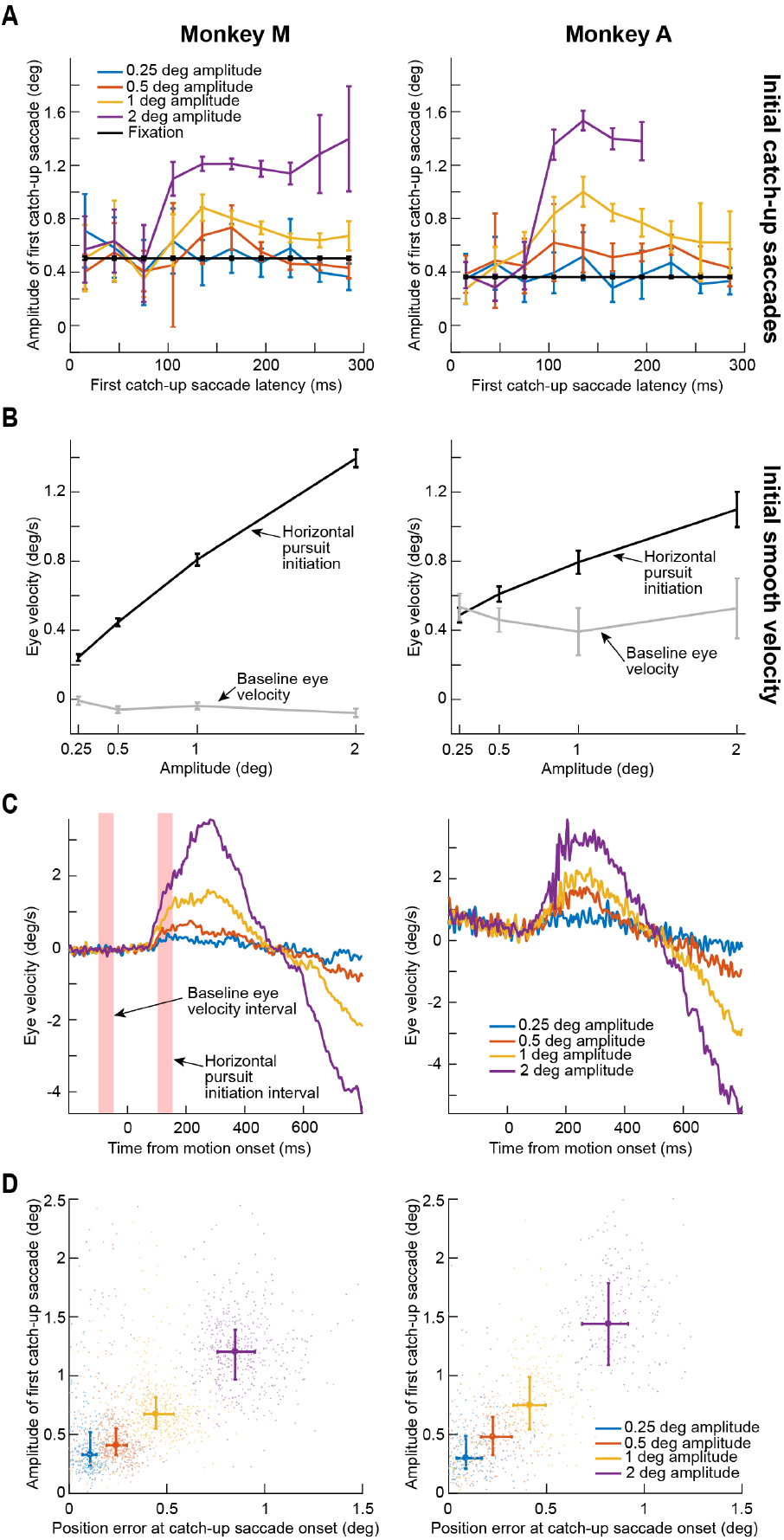
Interactions between saccades and smooth pursuit eye movements during initiation of tracking. (**A**) Amplitude of the first catch-up saccade after target motion onset as a function of time. Error bars denote 95% confidence intervals, and the different colors show different target motion amplitudes. The black curve shows microsaccade amplitude during steady-state fixation, for reference. For larger amplitude trajectories, first catch-up saccade amplitude increased with time, consistent with an expected increase in position error (see **D**). (**B**) A lack of (or weak) increase in first catch-up saccade amplitude in **A** (e.g. for 0.5 deg target motion amplitudes) might be because saccade-free smooth pursuit initiation may have already acted to reduce eye position error. For example, initial eye velocity during saccade-free initiation (see pink measurement intervals in the left panel of **C**) increased with increasing target amplitude. For monkey **A**, there was a small, systematic rightward drift in fixation, explaining this monkey’s gray curve. Also, note that in **A**, saccades with latencies less than ~50-60 ms were likely not initial catch-up saccades but instead fixational microsaccades. That is why their amplitudes were low for all conditions. Error bars denote 95% confidence intervals. (**C**) Example average saccade-free smooth velocity traces from horizontal tracking (like **B**) showing the measurement intervals for **B** and also the idea that smooth velocity effects increased with increasing target amplitude. (**D**) For each first catch-up saccade and target motion amplitude, we plotted saccade amplitude as a function of position error existing at saccade onset. Each dot shows an individual saccade, color-coded by the specific target amplitude trajectory in the experiment. The saturated, bold dots and error bars denote median and inter-quartile ranges (first to third quartile of the data).

Of course, the results of Fig. 10A do not necessarily describe individual movements and how they may have been affected by both position error and initial smooth velocity acceleration (occurring prior to saccade triggering). We therefore plotted, for each individual saccade, the position error that existed at the saccade onset. We found that the initial catch-up saccade amplitude increased with increasing position error for the same data as in Fig. 10A (Fig. 10D), and this analysis was indeed more sensitive than that in Fig. 10A. Naturally, regardless of the position error that existed at saccade onset, since the target was continuously moving, it was expected that the saccades, individually, would be additionally affected by target motion, as is known to happen (Fleuriet et al. 2011; Quinet and Goffart 2015). In the present study, we did not explicitly compare microsaccades during fixation to small catch-up saccades after initial target motion, but we expect similar results to (Fleuriet et al. 2011; Quinet and Goffart 2015).

Finally, we checked whether the first catch-up saccade latency itself depended on target amplitude. In Experiment 2, we plotted histograms of first catch-up saccade latency in the different amplitude conditions (Fig. 11). For the smallest amplitude trajectories, catch-up saccade latencies were long and variable. As target amplitude increased, latency became less variable, as well as shorter. Since Fig. 10 showed that it was more likely for catch-up saccade amplitudes to increase with increasing target position amplitude, these results constitute a monkey replication of human studies, showing that saccade latencies are substantially longer for very small-amplitude visually-guided saccades compared to larger ones (Kalesnykas and Hallett 1994).

**Figure 11.**
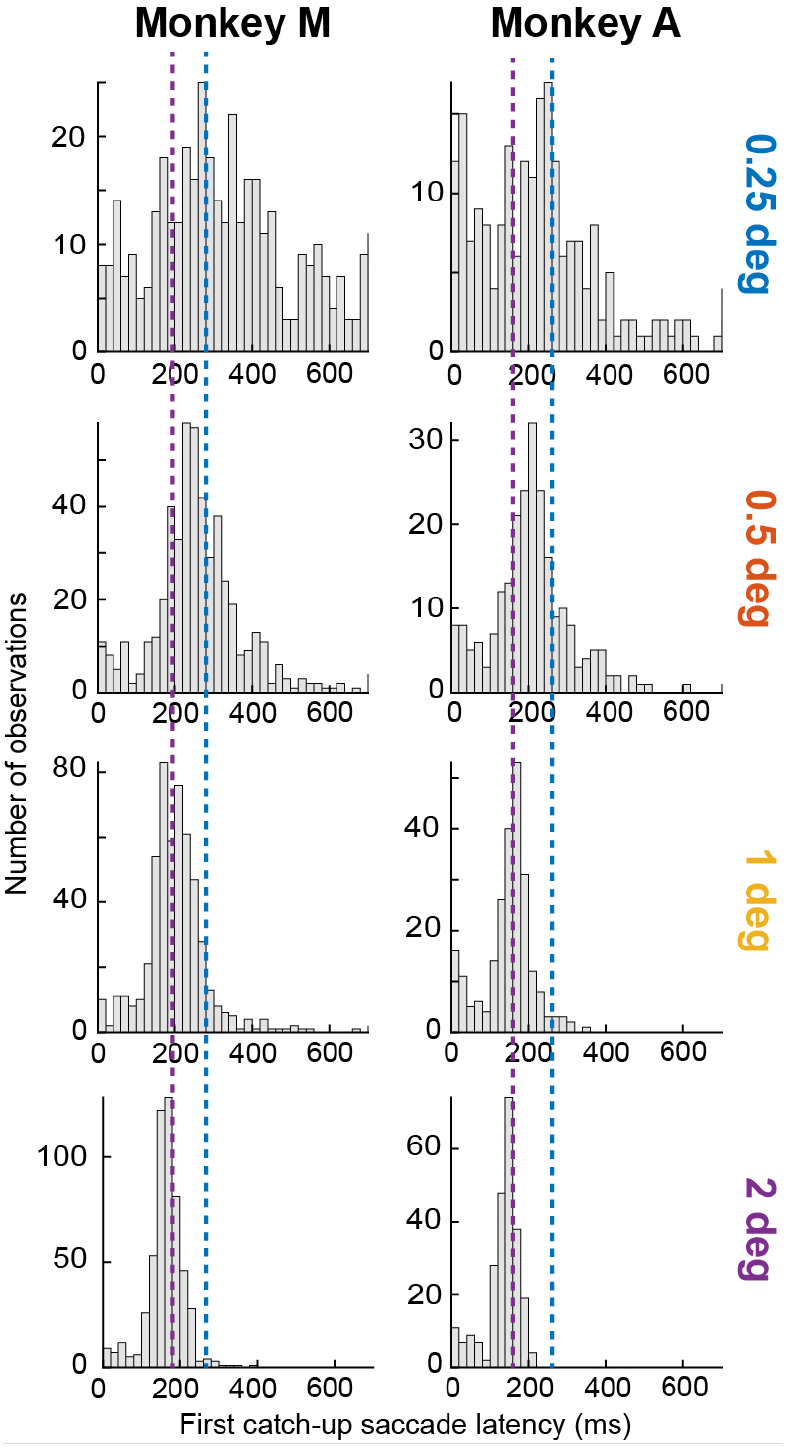
Dependence of first catch-up saccade latency on target amplitude. We plotted histograms of first catch-up saccade latency for different target amplitudes in Experiment 2. We show results only from horizontal tracking for clarity; other directions showed similar effects. Since larger target amplitudes resulted in larger catch-up saccades (e.g. Fig. 10), the present histograms show monkey replication of human observations that very small visually-guided saccades are associated with longer reaction times than larger ones (Kalesnykas and Hallett 1994). For example, for each monkey, the purple dashed line marks the latency bin with most observations at 2 deg target motion amplitude (bottom panel), and the blue dashed line marks the latency bin with the most observations at 0.25 deg target motion amplitude (top panel). As can be seen, there was a substantial differential in saccade times for the different target motion amplitudes. Note that, like in Fig. 10, the small distribution of saccades occurring with latencies <100 ms in this figure are likely not genuine first catch-up saccades, but instead fixational microsaccades. This is further supported by the noticeable dip in the histograms at ~100 ms, which is similar to the phenomenon of microsaccadic inhibition reported in the literature for both humans and monkeys (Buonocore et al. 2017; Hafed and Ignashchenkova 2013; Rolfs et al. 2008).

## Discussion

We attempted to characterize monkey oculomotor behavior with small-amplitude slow motion trajectories. We catalogued both smooth velocity effects as well as catch-up saccade effects. For smooth velocity effects, we found that pursuit gain was low for both low and high temporal frequencies, only reaching a peak in mid-frequencies near 1 Hz. This is in contrast to previous human experiments with large-amplitude (Collewijn and Tamminga 1984; Fabisch et al. 2009; Rottach et al. 1996) or small-amplitude (Martins et al. 1985) sinusoidal motions, in which low-pass behavior was observed. In terms of catch-up saccades, we found that they increased in frequency when smooth velocity gain was high, and they acted to increase the effective bandwidth of the overall tracking behavior up to 2 Hz. Moreover, we found that catch-up saccades during presentation of low temporal frequencies acted more like fixational microsaccades, whereas catch-up saccades during presentation of high temporal frequencies were large and infrequent.

Our results provide a necessary foundation for exploring the neural mechanisms subserving fixational ocular drifts in awake monkeys. This complements early characterizations of awake monkey smooth pursuit eye movements with higher amplitudes/speeds (Lisberger and Westbrook 1985). These early characterizations were themselves a major boon for a wide range of significant and seminal subsequent discoveries about the neural mechanisms for oculomotor control in general, and about the neural mechanisms for smooth pursuit in particular (Krauzlis 2004). Our next goal is to extend our current results by uncovering neural substrates in the same animals. While our primary interest lies in oculomotor control circuitry, it would also be interesting to relate small eye movements, like those that we have studied here, to the sensory drive associated with slow visual motion itself. For example, it is interesting to note that speed tuning preferences in motion-sensitive cortical area MT are overwhelmingly >1 deg/s (DeAngelis and Uka 2003; Inagaki et al. 2016). This might provide a neural constraint on smooth velocity gain at slow speeds, and it also leads to intriguing questions about how slow moving stimuli can be processed for ocular drift in general.

To support this future work, we were careful to avoid unnecessarily penalizing the monkeys for making saccades during tracking. Specifically, we aimed to minimize over-training on one particular movement modality. For example, early human studies with small-amplitude motions barely had any saccades in the experiments, to focus almost solely on slow control effects (Martins et al. 1985). However, we wanted the animals to engage in as naturalistic a behavior as possible, such that we could understand important interactions between slow control and micro/saccadic control. This has allowed us to make the interesting observation that smooth pursuit gain in our monkeys exhibited band-pass behavior, unlike in (Martins et al. 1985). This has also allowed us to demonstrate that there were actually more saccades when pursuit gain was high than when it was low.

Such an observation of a concomitant increase in catch-up saccade frequency along with an increased velocity gain might suggest that catch-up saccades normally behave like fixational microsaccades. The latter eye movements continuously re-align gaze with a foveal target under a variety of stimulus conditions (Guerrasio et al. 2010; Ko et al. 2010), and even when competing peripheral stimuli are presented (Tian et al. 2018; 2016). This means that their frequency of occurrence might, for instance, increase when the target is sharp and providing a clear spatial reference frame for re-aligning gaze. During pursuit, similar re-alignment of gaze is necessary, and the sharpness of the spot being tracked may be sufficient to increase the production of catch-up saccades. This is consistent with observations that foveal targets increase catch-up saccade frequency in humans (Heinen et al. 2018; Heinen et al. 2016); presumably, foveal targets not only support good smooth velocity gain, but they also provide the oculomotor system with a punctate spatial reference point to which gaze can be re-directed. We also found in monkeys and with larger-amplitude sinusoidal pursuit that there was a tendency for higher catch-up saccade frequencies for smaller foveal pursuit targets rather than for bigger and fuzzier ones (Hafed et al. 2008). It would be interesting to analyze the relationships between eye position error and catch-up saccade likelihood with a foveal target in more detail, such that one can uncover an almost-deterministic estimate of whether a catch-up saccade can occur at any one moment of time or not, along the lines of (de Brouwer et al. 2002). This kind of approach was recently made for microsaccades (Tian et al. 2018), and it is very intriguing because predicting whether and when a microsaccade might take place can, at least in principle, be used to estimate the occurrence of distinct cognitive performance effects associated with such movements (Bellet et al. 2017; Chen et al. 2015; Hafed 2013).

Another interesting aspect of catch-up saccades in our study was related to how they exhibited complementary roles for retinal slip and position error at different frequencies. While catch-up saccades corrected for position error at low temporal frequencies (Figs. 4, 5), their biggest impact on position error occurred when smooth velocity gain declined sharply at 2 Hz. On the other hand, the impact of these saccades on retinal slip was overwhelmingly at low temporal frequencies (Fig. 5). This could be a function of the range of position and velocity errors that the oculomotor system had to encounter at the different temporal frequencies in our experiments.

Our interest in relying on more naturalistic tracking (i.e. with combined smooth and saccadic eye movements) may also explain why we observed band-pass smooth pursuit gain effects in our monkeys, even though a similar human experiment with low frequencies and small-amplitude trajectories found very high gain (Martins et al. 1985). As stated earlier, in that study, the human subjects tested were thoroughly trained, and they were instructed to minimize saccade generation. As a result, substantial systematic eye position drifts occurred in their experiments, whereas we did not observe such systematic drifts. In our case, we relied more on the natural behavior of the monkeys in being intrinsically interested to foveate the small white spot that was presented on the display. Yes, the monkeys became highly trained in the lab after multiple sessions, but their behavior was not experimentally shaped, say, by aborting trials whenever a saccade occurred. Instead, we rewarded them for tracking the target to within a reasonable radius, which may not be too different from natural variability in human fixation among untrained individuals (Cherici et al. 2012). As a result, our monkeys tracked the target with both smooth and saccadic eye movements.

In any case, our results provide complementary data to the pioneering work of Martins et al. (1985) using similar paradigms in humans. Our results are also in line with Cunitz’s (1970) interpretations about slow ocular drifts. However, one question that may arise from our experiments, and those earlier human studies, is concerning whether smooth pursuit is indeed analogous to slow ocular drifts or not (Cunitz, 1970; Martins et al., 1985; Nachmias, 1961). Specifically, the relatively low gain that we observed at low speeds/temporal frequencies may be interpreted as revealing a qualitative difference from a system that uses slow motor control of ocular drifts. In other words, it may be the case that there are two control systems, one for compensating for (noisy) ocular drifts during fixation and one for tracking moving targets. This remains to be seen, since an alternative interpretation is that the oculomotor system engages in different regimes of tolerance for position and velocity errors at different scales of eye movement (Cunitz, 1970). At the very slow speeds associated with the lowest temporal frequencies (e.g. 0.2 Hz), the retinal image motions experienced by the oculomotor system may be within its tolerance range for eye position control, and this would also be true for a stationary fixation stimulus. It would be interesting to identify visual and/or cognitive conditions in which it would be advantageous for the monkeys to increase their smooth velocity gain at low temporal frequencies, perhaps through reward or the use of challenging visual discriminations. If the monkeys do indeed increase their smooth velocity gain, this would also reconcile with the human results of (Martins et al., 1985).

We were also intrigued by our oblique tracking effects in Experiment 2. Catch-up saccade amplitudes increased in oblique tracking relative to cardinal-direction tracking. Moreover, the velocity gain for the horizontal component of oblique tracking was consistently higher than the velocity gain for the vertical component. The catch-up saccade amplitude increase likely reflected the slightly faster trajectories associated with oblique target motions in our stimulus design relative to cardinal target motions (Materials and Methods). As for the asymmetry between horizontal and vertical components of smooth velocity gain, this could relate to oblique effects in both motion perception and smooth pursuit (Krukowski and Stone 2005), and it is consistent with large-amplitude pursuit effects in humans (Rottach et al. 1996). We find the increase in horizontal versus vertical component of smooth velocity gain in our particular scenario of small-amplitude tracking additionally intriguing because it might relate to observations that small visually-guided saccades in humans overshoot their targets (Kalesnykas and Hallett 1994). We may have thus observed a similar phenomenon for smooth pursuit, at least for the horizontal component. In other words, it may be the case that small-amplitude smooth pursuit overshoots targets like small-amplitude saccades overshoot targets. This might add, at least in a correlative way, to evidence that smooth pursuit and saccades share neural resources (Krauzlis and Dill 2002; Krauzlis 2004; Krauzlis et al. 1997; Krauzlis et al. 2017). Similarly, even our catch-up saccade effects of Fig. 11 demonstrate a monkey correlate of human observations that visually-guided saccade latency increases for small target eccentricities. It would be interesting to extend these effects for other types of saccades, like delayed visually- or memory-guided saccades. In all, our results add to an extensive cataloguing in the literature of rhesus macaque sensory, perceptual, cognitive, and motor capabilities, testifying to the tremendous value of such an animal model for systems neuroscience research.

## Acknowledgments

We were funded by the Werner Reichardt Centre for Integrative Neuroscience (CIN) at the Eberhard Karls University of Tübingen. The CIN is an Excellence Cluster funded by the Deutsche Forschungsgemeinschaft (DFG) within the framework of the Excellence Initiative (EXC 307). We were also funded by the DFG through a Research Unit (FOR 1847). ZMH was additionally funded by the Hertie Institute for Clinical Brain Research at the Eberhard Karls University of Tübingen.

